# Breeding of microbiomes conferring salt tolerance to plants

**DOI:** 10.1101/2025.01.13.632835

**Authors:** Caio Guilherme Pereira, Joseph A. Edwards, Albina Khasanova, Alexis Carlson, Vanessa Brisson, Estelle Schaefer, Tijana Glavina del Rio, Susannah Tringe, John P. Vogel, David L. Des Marais, Thomas E. Juenger, Ulrich G. Mueller

**Affiliations:** Department of Integrative Biology, The University of Texas at Austin, Austin, TX, USA; Department of Plant Pathology and Microbiology, Texas A&M University, College Station, TX, USA; U.S. Department of Energy Joint Genome Institute, Lawrence Berkeley National Laboratory, Berkeley, CA, USA; Department of Civil and Environmental Engineering, Massachusetts Institute of Technology, Cambridge, MA, USA

**Keywords:** Microbiome breeding, host-mediated selection, plant-microbe interactions, *Brachypodium distachyon*, abiotic stress, sodium stress, aluminium stress, microbial co-occurrence networks, plant growth-promoting bacteria, sustainable agriculture

## Abstract

Microbiome breeding by host-mediated selection is a technique to artificially select for microbiomes conferring benefits to plants. Here, we describe leaf ionomics, microbial community composition, and network analyses of a microbiome-breeding experiment to generate microbiomes conferring salt tolerance to *Brachypodium distachyon*, a model for cereal crops. Plants receiving microbiomes selected to confer tolerance to either sodium- or aluminium-stress produced 69-198% higher total seed weight than plants receiving control microbiomes. Sodium-selected microbiomes reduced leaf-sodium concentration by 50%, whereas aluminium-selected microbiomes had no effect on leaf-tissue nutrient concentration, suggesting different mechanisms underlying microbiome-mediated salt tolerance. By testing these selected microbiomes in a cross-fostering experiment, we show that artificially-selected microbiomes attain (a) ecological robustness contributing to transplantability (inheritance) of microbiome-encoded effects between plants; and (b) network features identifying key bacteria promoting stress tolerance. Combined, these findings elucidate critical mechanisms underlying host-mediated selection as a tool to breed beneficial microbiomes in an agricultural context.

---

Root-associated microbial communities play fundamental roles in plant growth and development, affecting diverse aspects of the plant’s physiology^1^ and modulating its systemic defenses^2^, disease resistance^3^, and tolerance to both biotic and abiotic stresses^4,5^. As such, there has been an increasing interest in using plant-growth-promoting bacteria – or even entire microbial communities – in agricultural settings to increase yield and attenuate environmental stress. However, beyond the difficulty in identifying and isolating the key microbial players in a given microbiome, reliable bacterial inoculation of plants is often hindered by differences in soil properties between the source and novel environment, as well as by competition with the native microbiota^6^. To address these challenges, two approaches have been proposed: a bottom-up approach involves designing synthetic microbial communities from first principles^7–9^, whilst a top-down approach involves microbiome breeding through host-mediated selection, a flexible framework to artificially select for microbiomes with desired properties to plants.

Microbiome breeding through host-mediated selection is a process in which microbiome functions can be artificially selected using host-plant performance as proxy to gauge microbiome properties, whilst remaining agnostic to the microbial community composition and the ecological and evolutionary forces that ultimately shaped it^10,11^. By harvesting microbiomes from plants with desired phenotypes (e.g., tolerance to salt stress) and using these microbiomes to inoculate subsequent cohorts of plants, it is possible to steer microbiome composition - and therefore microbiome function - towards helping plants cope with said stress^11,12^. A major advantage of this method is that it leverages properties of the whole microbiome, including synergistic and emergent properties that would be difficult to predicted from first principles^13^. It is also much more likely to produce ecologically stable, transplantable communities. Different microbiome breeding techniques have been used to successfully enhance growth or flowering in model species and crops, including *Arabidopsis*^14,15^ and wheat^16^, though with highly variable and overall modest results.

We recently developed a microbiome selection protocol (Fig. 1) to generate microbiomes conferring salt tolerance to *Brachypodium distachyon*^17^, a model for cereal crops and grasses^18,19^. Our methods aim to optimize microbiome breeding through manipulations that: 1) Increase plant control over their microbiome; 2) maximize the transplantability (inheritance) of microbiome properties; and 3) minimize the impact of migration and drift throughout the microbiome selection process. With these manipulations, we were able to increase plant productivity of plants experiencing sodium and aluminium stress after only 3 rounds of selection^17^, and reported preliminary results regarding the effects of our selected microbiomes on seed production of plants under extreme salt stress^17^. Here, we re-analyse these data in combination with physiological evidence, showing with bioinformatic analyses and plant-tissue ionomics that our host-mediated artificial-selection protocol generated microbiomes with unprecedented effects on plant fitness, and proposing mechanisms underlying the microbiome-mediated influence on osmotic and heavy-metal stress tolerance in plants. More importantly, we expand on that study by presenting the community composition analyses of the microbiomes generated in our breeding experiments, showing that both the composition and interaction patterns between microbes changed depending on the salt stress in which microbiomes were selected upon (referred hereafter as ‘selection history’) and on the stress they currently experience.

**Figure 1.**
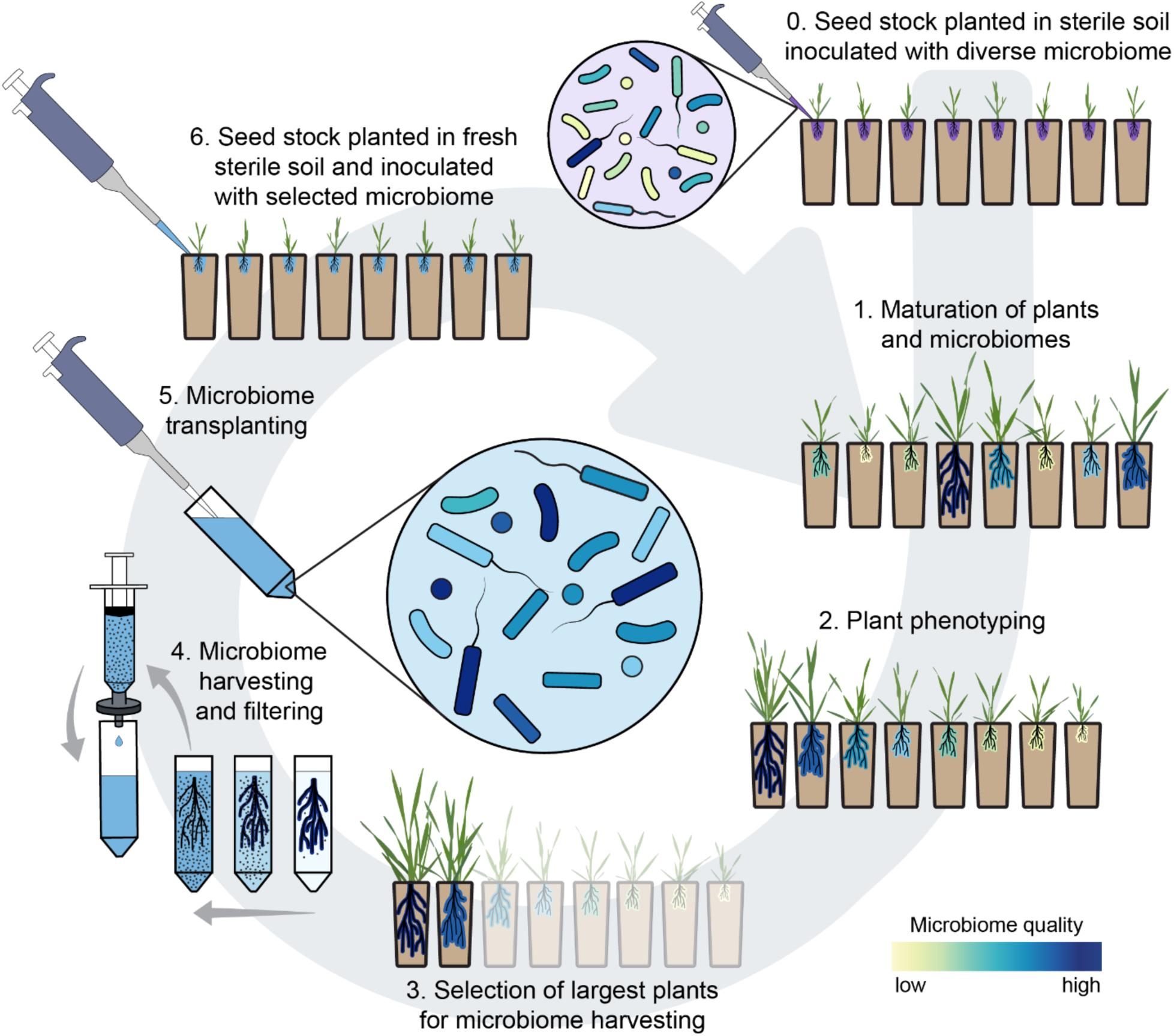
Host-mediated artificial-selection scheme used for microbiome breeding. Seedlings from a non-evolving seedstock were inoculated with a taxonomically diverse, initial microbiome composed of root-associated bacteria collected from several greenhouse-grown *Brachypodium distachyon* plants and other grass species collected from the field (Step 0). Plants and associated microbiomes were allowed to mature and interact for ∼28 days (Step 1), then plants were phenotyped according to size (Step 2). The two largest plants of each selection line were selected for microbiome harvest, which consisted of submerging the roots in extraction buffer and gently shaking them until most material adhering to the root was released into the solution buffer, which was then filtered to remove soil particles and larger-sized organisms that otherwise would co-propagate with the bacterial fraction (Steps 3-4). This bacterial microbiome fraction was then inoculated into a fresh batch of seedlings from the same non-evolving *B. distachyon* seedstock (Steps 5-6), planted in the same growth media, but now stressed with a slightly higher concentration of either aluminium or sodium sulphate (see Supplementary Table S2 for salt concentrations used throughout the experiment). The selection cycle depicted between steps 1-6 illustrates a single round of selection, for a single selection line (each composed of 8 plants per microbiome-generation), and cycles were repeated 9x throughout our experiment.

## Results

After nine rounds of microbiome selection (Fig. 1), where salt stress was incrementally increased in each microbiome generation, we tested the effect of our selected microbiomes on plant fitness in a cross-fostering experiment (Fig. 2a). Plants that received selected microbiomes showed, on average, 69-198% higher total seed weight than plants inoculated with fallow-soil control microbiomes (Fig. 2b), suggesting that the host-mediated selection generated microbial communities that improve plant fitness (expressed as seed productivity) under salt stress. This growth-promotion can be attributed to bacterial components within the selected microbiomes, given that solute controls (filtered to remove bacteria) did not affect seed output to the same extent: they increased total seed weight by <30%, and only under aluminium stress (Fig. 2b). Interestingly, sodium-selected microbiomes increased fitness of plants experiencing aluminium stress and *vice-versa*, suggesting a partial overlap in some functional property of these selected microbiomes. As expected, plants showed highest seed output when inoculated with the microbiome selected under the same stress they were experiencing in the cross-fostering experiment (sodium-selected microbiomes for sodium-stressed plants, and aluminium-selected microbiomes for aluminium-stressed plants; Fig. 2b).

**Figure 2.**
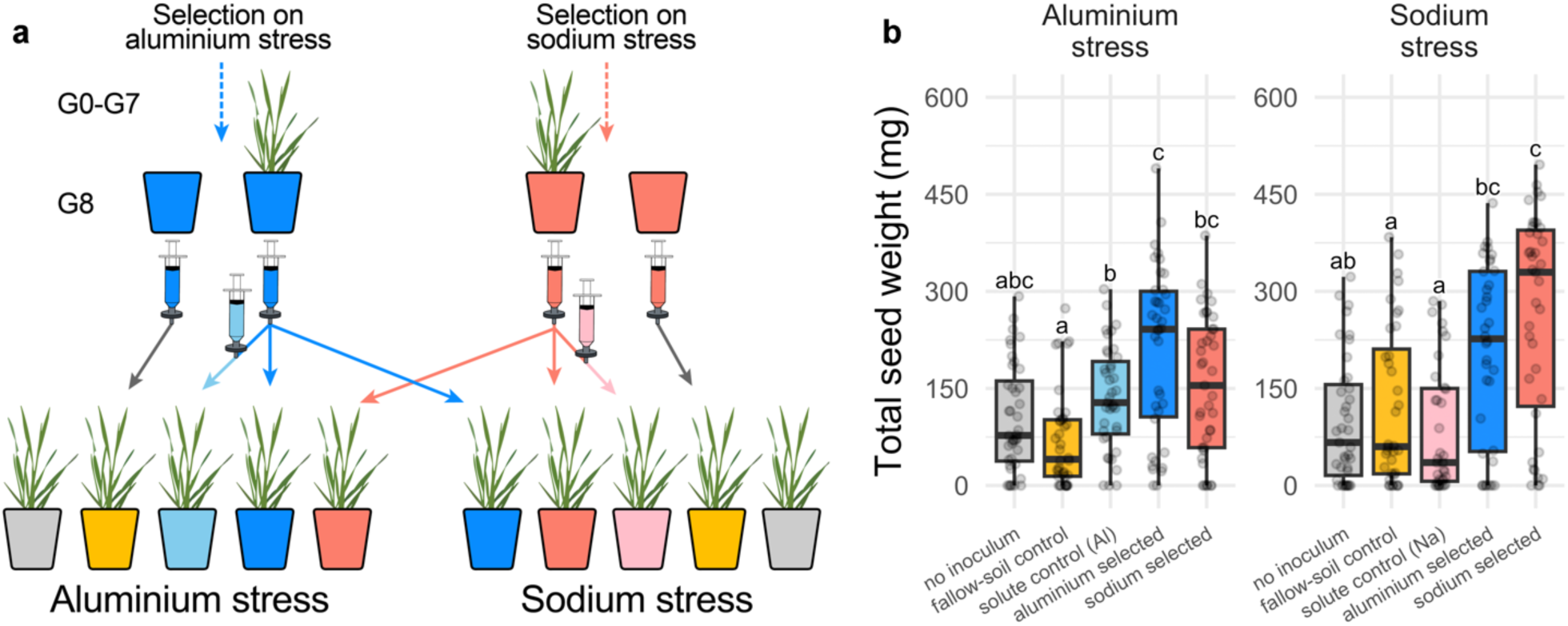
Experimental design for microbiome inoculation at microbiome-generation 9 and its effect on plant fitness. **(a)** Plants from the same seedstock as the ones used in selection cycles (Fig. 1) were grown under aluminium or sodium stress for ∼10 weeks after receiving one of the following: 1) aluminium-selected microbiomes that were 2µm-filtered to propagate bacteria (blue); 2) sodium-selected microbiomes that were 2µm-filtered to propagate bacteria (red); 3) double-filtered solutes from aluminium- or sodium-selected microbiomes, where selected microbiomes were passed through both 2µm and 0.2μm filters to remove live bacterial cells, so any effect of this filtrate on plant fitness would be the result of root exudates or viruses that may have passed through the 0.2μm filter (light blue or light pink, respectively); 4) fallow-soil microbiomes, where bacterial microbiomes were harvested from root-free fallow soil subject to the same ramping of the imposed stresses through nine generations (yellow); 5) no inoculum (grey). The experiment was conducted using a partial “cross-fostering” design, where both 2µm-filtered aluminium-selected and 2µm-filtered sodium-selected microbiomes were tested on plants under each salt stress, whereas the double-filtered aluminium-selected solutes were only tested on plants under aluminium stress, and the double-filtered sodium-selected solutes were only tested on plants under sodium stress. **(b)** Plant fitness, expressed as total seed weight, of plants given different inocula at generation 9. Boxplots show medians, 25^th^, and 75^th^ percentiles. Whiskers extend to 1.5 times the interquartile range and data points presented beyond whiskers represent outliers. Different letters indicate significant differences between bacterial treatments within individual salt stresses based on Šídák posthoc tests.

### Selected microbiomes affect plant tissue-nutrient composition differently

To explore possible mechanisms through which the selected microbiomes helped plants cope with salt stress, we analysed the leaf-nutrient composition of plants grown with distinct microbiomes for signals of differential accumulation or exclusion. We found that the sodium-selected microbiomes led to a ∼50% decrease in leaf-sodium concentration of plants growing under sodium stress (Fig. 3a). This suggests that the sodium-selected microbiomes either reduce sodium availability in the soil, or more likely, elicit a plant response (e.g., increasing proline synthesis^20^, modulating phytohormone production^21^) to cope with the stress. We found no difference in leaf-aluminium concentration among plants that received our selected microbiomes and those that received solute-control microbiomes (no live bacteria), fallow-soil microbiomes, or no microbial inoculum (Fig. 3a), regardless of the salt stress those plants were experiencing. This suggests that the fitness benefit that aluminium-selected microbiomes conferred to plants under aluminium stress (Fig. 2b) was not the result of a reduction in aluminium uptake or accumulation.

**Figure 3.**
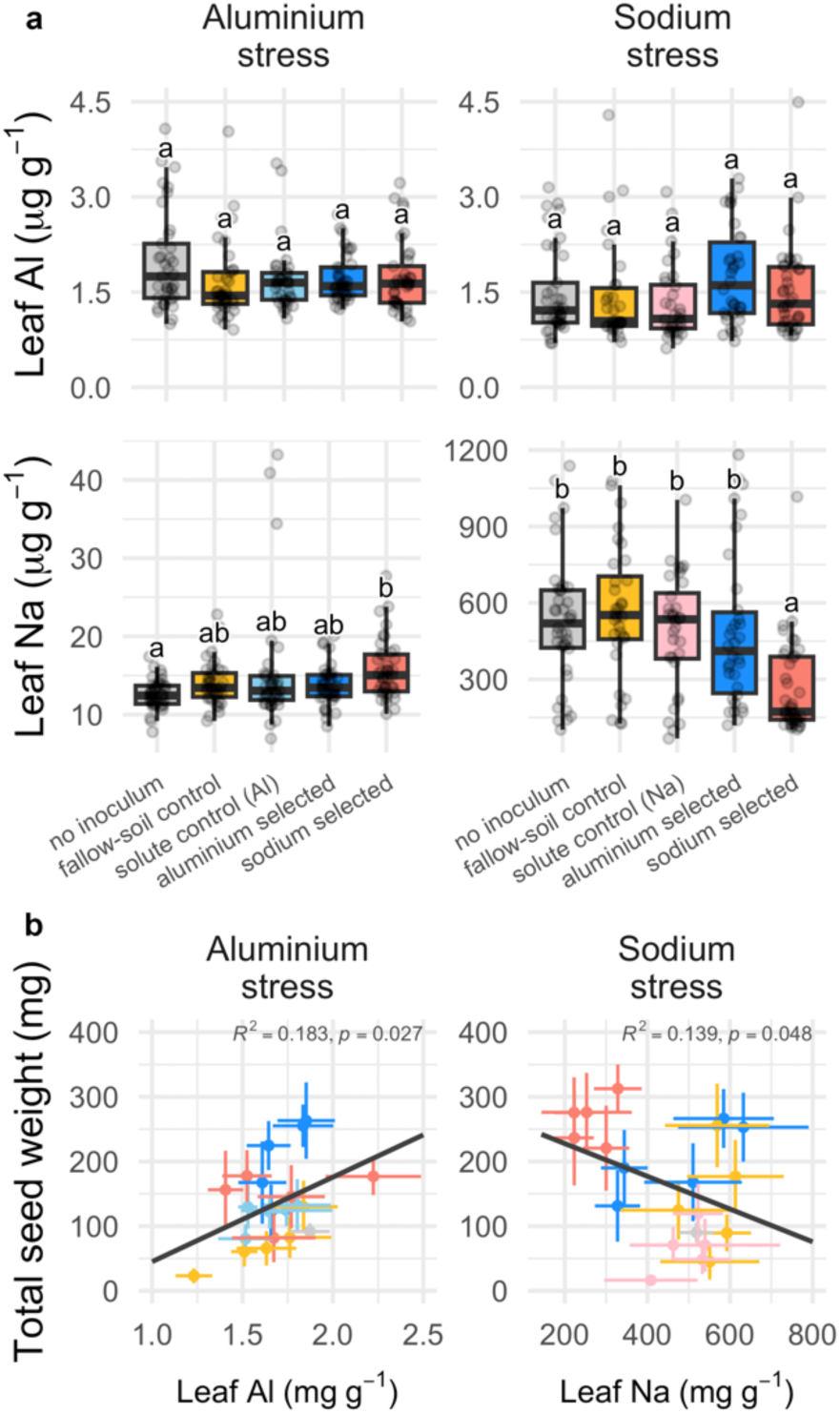
Leaf aluminium (Al) and sodium (Na) concentration of plants given different microbiome inocula at generation 9 and their effect on plant fitness. **(a)** Boxplots of leaf-tissue Al and Na concentrations at both salt stresses, with medians, 25^th^, and 75^th^ percentiles. Whiskers extend to 1.5 times the interquartile range, and data points beyond whiskers represent outliers. Different letters indicate significant differences between bacterial treatments within individual salt stresses based on Šídák posthoc tests. (**b**) Correlations between plant fitness (expressed as total seed weight) and leaf Al and Na concentrations. Individual data points represent mean values for each selection line, so error bars are depicted along both axes. Regression lines are depicted in black, with *R*^2^ and *p*-values annotated above. The colours match the microbiome treatments illustrated in panels (a), with grey representing ‘no inoculum’, yellow representing ‘fallow-soil control’, light blue and pink representing ‘solute controls’ (aluminium- and sodium-selected microbiomes, respectively), blue representing ‘aluminium-selected microbiome’, and red representing ‘sodium-selected microbiome’.

The correlation between seed weight and leaf-sodium concentration (Fig. 3b) suggests that sodium exclusion was the likely mechanism underlying the sodium-selected microbiomes’ positive impact on plants under sodium stress, with seed output decreasing with increasing leaf-sodium concentration (*R*^2^=0.139, *p*=0.048) and plants that received sodium-selected microbiomes clustering in the “low-sodium/high-fitness” portion of the graph (Fig. 3b). In terms of the relationship between seed productivity and leaf-aluminium concentration, there was a positive correlation (*R*^2^=0.183, *p*=0.027; Fig. 3b) but this was driven by a couple of selection lines. Still, whether significant or not, the correlation pattern does not support an aluminium-exclusion hypothesis, which predicts a negative correlation.

### Microbiome breeding generates stress-specific bacterial communities robust to perturbation

Next, we assessed how microbiome breeding affected root-associated bacterial community composition. Because the effectiveness of our selected microbiomes was strongly dependent on their selection history (Fig. 2b), we hypothesized that our sodium- and aluminium-selected microbiomes – despite being derived from the same initial inoculum – would diverge in composition over multiple rounds of selection. As expected, plants that received inoculum from selected microbiome lines hosted communities distinct from null-inoculated plants (i.e., plants whose microbiomes “rained in” entirely from the growth-chamber environment; Fig. 4a). Plants inoculated with selected microbiomes also acquired distinct root-associated bacterial communities: we found two distinct clusters corresponding to the two selection histories in a principal coordinate analysis (Fig. 4b). Selection history explained the largest amount of variance of assembled community composition (*R*^2^=0.38, *p*<0.001), followed by salt stress (*R*^2^=0.06, *p*<0.001). There was also a significant interaction between selection history and salt stress on microbiome composition (*R*^2^=0.08, *p*<0.001), indicating that the effect of selection history on microbiome composition was contingent on the salt stress plants were experiencing. The sodium-selected microbiomes showed greater “plasticity” (referred here as dispersion in PCoA plots) under both sodium and aluminium stress than aluminium-selected microbiomes (Fig. 4a-c), likely due to their higher compositional variability and Shannon diversity (Supplementary Fig. S1). Together, these results indicate that: 1) the community composition of our selected microbiomes are ecologically robust when exposed to a new salt stress (Fig. 4c); 2.) functionally important microbiome-components can be perpetuated with sufficient fidelity across microbiome generations; and 3.) the microbiome-mediated robustness to salt stress observed in this study was not due to environmental microbes that were secondarily acquired during the cross-fostering experiment.

**Figure 4.**
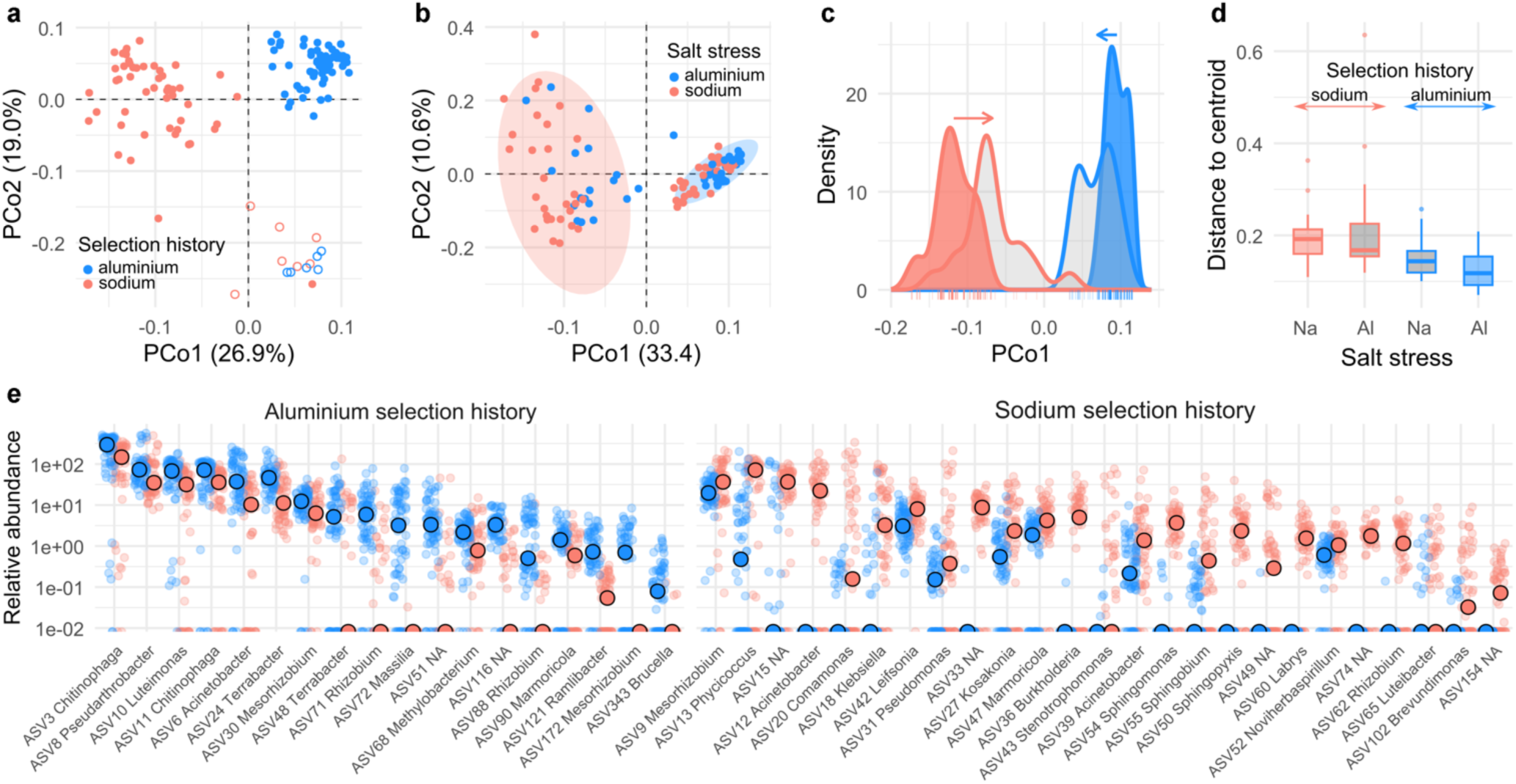
Microbiome breeding generates divergent and ecologically robust bacterial communities. **(a)** PCoA of root-associated bacterial communities from the 2×2 cross-fostering experiment after nine rounds of selection; closed dots represent samples receiving inoculum from the previous generation while open points are samples from plants receiving no-inoculum (i.e., colonized by environmentally derived bacteria only). **(b)** PCoA of the same experiment but omitting null-inoculated samples; background ellipses depict ‘selection history’, whilst point colour represents the ‘salt stress’ each host plant (and microbiome) experienced in the experiment. **(c)** Density-distribution of samples along PCo1 of panel **b**.; the outline colour of the distribution curves depicts the ‘selection history’ prior to microbiome-generation 9, whilst the fill colour represents the ‘salt stress’ experienced by plants of microbiome-generation 9, with grey fill illustrating the distribution of samples experiencing a salt stress alternative to their selection history. **(d)** Dispersion of datapoints around their centroid indicates that sodium-selected microbiomes are more variable in their composition than aluminium-selected microbiomes. **(e)** ASVs identified as differentially abundant between the two selection histories; large points represent the median relative abundance of the ASV in each ‘salt stress’ treatment, with the clouds of small points depicting the relative abundances for all samples.

To identify differentially abundant bacterial taxa across selection histories, we modelled the relative abundance of amplicon sequence variants (ASVs) detectable in >25% of the samples. Of the 62 ASVs meeting this prevalence threshold, 43 were differentially abundant depending upon the selection history (adjusted *p*<0.01, Fig. 4e): 25 ASVs showed significantly greater relative abundance in sodium-selected microbiomes and 18 in aluminium-selected microbiomes. The number of total ASVs included in these models was too low to identify broad taxonomic enrichments, yet there were clades enriched in selected microbiomes: e.g., several ASVs from the Chitinophagales order were enriched in aluminium-selected microbiomes, whilst ASVs from the Cyanobacteria phylum and Sphingomonadales order were enriched in the sodium-selected lines. Most microbes enriched in one selection background had a median abundance of zero in the alternative background; this was the case for 17 out of the 25 ASVs significantly enriched in the sodium-selection history and 8 out of the 18 ASVs enriched in the aluminium-selection history. This suggest that the mechanism underlying the aforementioned robustness of the microbiome composition in the face of a different stress is associated with diversity loss, and that through purifying selection or drift, some microbes became extinct, thus limiting the degree to which microbiome composition can adjust to an alternative stress.

### Microbiome breeding shapes microbial co-occurrence network structure and topology

To test the effect of microbiome breeding on microbial community structure and topology, including network complexity and stability, we estimated microbial co-occurrence networks^22–24^ for plants given null-, sodium-, and aluminium-selected microbiomes under either sodium or aluminium stress. Limiting each network to 50 nodes, we found a total of 113 unique ASVs from 32 bacterial families (Fig. 5a) across the six inferred networks (Fig. 5b). Despite the high alpha-diversity recovered from plants given null and selected microbiomes (781 ASVs across all treatments), these 113 ASVs represented >98.1% of the total sequencing reads of each library (Table 1).

**Figure 5.**
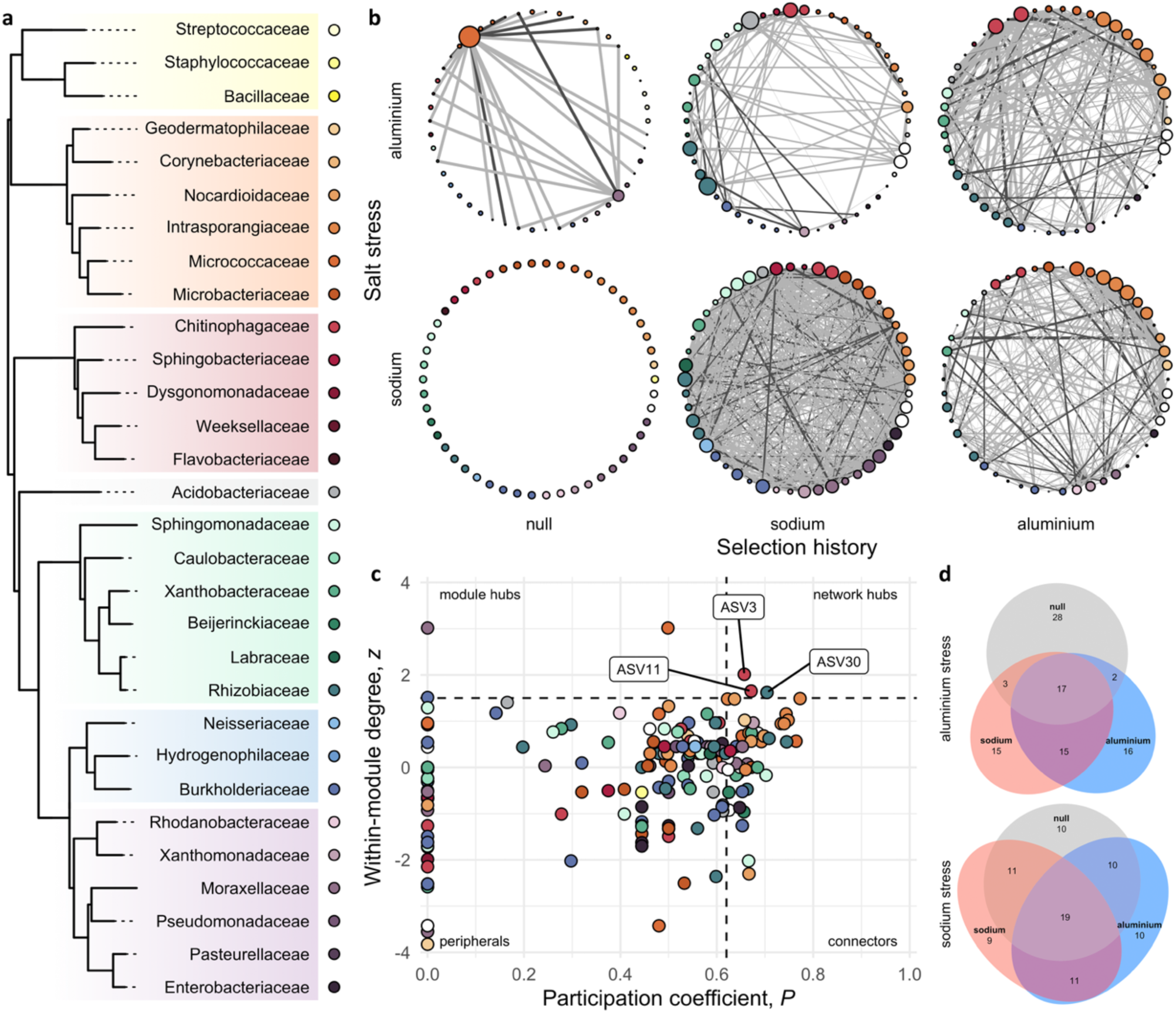
Microbial co-occurrence networks for root-associated bacteria after 9 generations of artificial selection. **(a)** Phylogenetic relationship among bacterial families of ASVs identified in our network analyses, as extracted from the maximum-likelihood estimated molecular phylogeny of life^66^; each bacterial family is represented by a unique colour, with different hues depicting classes: yellow for Bacilli, orange for Actinobacteria, red for Bacteroidia, grey for Acidobacteriia, green for α-Proteobacteria, blue for β -Proteobacteria, and purple for γ-Proteobacteria. **(b)** Microbial co-occurrence networks inferred for plants given null, sodium-selected, or aluminium-selected microbiomes under either sodium or aluminium stress. Node sizes are proportional to their degree in each graph, with unconnected nodes set a defined size to allow the identification of family/class (as illustrated in the phylogeny). Light grey edges represent positive correlations between nodes, dark grey edges represent negative correlations (line thickness is proportional to the magnitude of the correlation in either case). **(c)** *z*-*P* plot exhibiting patterns of within- and across-module connectivity of the ASVs from the six networks shown in ‘**b**’, with the identified network hubs highlighted. **(d)** Venn diagram showing the number of unique and shared ASVs from the network analyses depicted in ‘**b**’; set of ASVs identified only in plants that received null microbiomes is coloured in grey, whilst those that were found only in plants that received sodium- or aluminium-selected microbiomes are coloured in orange and blue, respectively.

**Table 1.** Network parameters of *Brachypodium distachyon* (Bd3-1) root-associated microbiomes at two salt stresses (aluminium or sodium sulphate) after 9 rounds of artificial selection for sodium and aluminium tolerance, as well as the unselected control (null) control. The “sodium stress - null selection history” network was empty, thus the absence of results. Percentage of reads presented as mean ± se.

| Stress | Selection history | Samples | Percentage of reads | Components | Clustering coefficient | Modularity | Positive edge percentage | Edge density | Natural connectivity |
| --- | --- | --- | --- | --- | --- | --- | --- | --- | --- |
| aluminium | null | 6 | 99.6 ± 0.08 | 29 | 0.00 | 0.19 | 41.18 | 0.02 | 0.05 |
|  | sodium | 29 | 98.1 ± 0.60 | 20 | 0.74 | 0.57 | 84.15 | 0.07 | 0.05 |
|  | aluminium | 30 | 99.9 ± 0.02 | 3 | 0.45 | 0.21 | 49.34 | 0.20 | 0.11 |
| sodium | null | 8 | 99.4 ± 0.11 | - | - | - | - | - | - |
|  | sodium | 37 | 98.5 ± 0.39 | 1 | 0.74 | 0.13 | 55.20 | 0.54 | 0.19 |
|  | aluminium | 30 | 99.8 ± 0.03 | 2 | 0.49 | 0.18 | 43.52 | 0.21 | 0.08 |

Both selection history and salt stress affected network topology, with sodium- and aluminium-selected microbiome networks showing a much higher edge density (percentage of all possible connections that are realized within a network) than the null (i.e., no inoculate) microbiome networks (Fig. 5b; Table 1). Interestingly, the edge density was highest when salt stress matched the microbiome selection history. The number of components followed an inverse trend, being lower in the selected microbiomes and lowest when salt stress matched selection history, likely because the higher number of edges helped “bridge the gap” between the largest connected components and small clusters and isolated nodes. The clustering coefficient, on the other hand, was strongly correlated with microbiome selection history, with little to no effect of current salt stress (Table 1); selected-microbiome networks had a higher clustering coefficient than null-selected ones, with sodium-selected lines showing the highest values. Together, these results suggest that salt stress and, more importantly, selection history led to an increase in network complexity. In addition, sodium- and aluminium-selected microbiomes showed highest natural connectivity (a measure of robustness of complex networks^25^) under sodium and aluminium salt stresses, respectively (Table 1; Supplementary Table S1).

Changes in network complexity could reflect changes in the roles of individual bacteria within their communities under different contexts of selection and stress: ASV3 and ASV11, for example, are two *Chitinophaga* ASVs that we found in the microbiome of all plants under aluminium stress, but only in aluminium-selected microbiomes did they show the pattern of within-module degree and participation coefficient that is consistent with a network hub (Fig. 5c; Supplementary Fig. S3).

This means that they may act as keystone species^26^, exerting a disproportional influence in the structure of the microbiome and its function under aluminium stress, but this is highly dependent on the selection history that shaped the community composition. In addition to individual nodes, there were also certain groups of bacteria that were enriched during selection and that, collectively, contributed disproportionally to the connectance of their network; aluminium-selected microbiomes, for example, had several members of Intrasporangiaceae, including species of *Janibacter* and *Terrabacter*, all with relatively high degrees. Interestingly, species from these two genera have been shown to be involved in biological phosphate removal from wastewater and bioremediation processes^27^. It is important to note, however, that whilst the selection history set the initial conditions of these microbiomes by differentially perpetuating microbes through multiple generations, the end result was the outcome of a complex interaction between past selection and current stress, as the composition (and likely structure and function) of the same microbiome can vary depending on the conditions into which they are inoculated (Fig. 5d).

## Discussion

Microbiome breeding as a method for shaping microbial communities with beneficial properties for plants has been around for years, yet progress in what some have described as the next-generation of breeding strategies^12^ has been slow, mainly due to the lack of studies testing its theoretical predictions and overall effectiveness. Here, we address this gap by describing the results of a host-mediated artificial-selection experiment, designed within a rigorous eco-evolutionary framework, to breed microbiomes conferring salt tolerance to plants. Guided by quantitative-genetic principles^28,10^, our protocol aimed to maximize the heritability of microbiome effects^29^ on plant traits, as well as the transmission fidelity and microbiome stability throughout the selection cycles, and resulted in selected microbiomes that increased plant fitness by up to 198% under salt stress^17^. We successfully shaped microbiomes to help plants cope with osmotic (sodium) and heavy-metal (aluminium) stress, and their effects on leaf-nutrient composition implicate two fundamentally different mechanisms underlying the microbiome-mediated salt tolerance of host plants. We show that microbiomes bred under different salt stresses became compositionally distinct and, by testing their effects in a cross-fostering experiment, ecologically robust to perturbation. Finally, we show that our selection protocol affected not only the composition of bacterial communities, but also their putative interaction patterns, increasing microbial co-occurrence network complexity and stability.

Previously, we showed that microbiome breeding through successive and differential propagation of microbiome components is not only possible, but potentially fast, with only 1-3 selections cycles needed to generate microbiomes that improve plant growth by >38%^17^. After nine cycles, we subjected our selected microbiomes through a partial cross-fostering experiment to test their effectiveness, where we found, in a previous analysis, an overall positive effect on seed output, with split results in terms of specificity^17^. Specifically, aluminium-selected microbiomes conferred tolerance to both aluminium- and sodium-stressed plants, whilst sodium-selected microbiomes only benefited plants under sodium stress. Here, we first re-analysed these results, incorporating ‘selection line’ as a factor and using ‘fallow-soil’ as control, given the unrealistic nature of a true ‘null control’ in real-life conditions and because it has been hypothesized that plants may sometimes - particularly during the earlier stages of growth - show increased vigor in sterile conditions due to either a complete lack of detrimental bacteria in the soil^30,31^ or an increased nutrient availability (i.e., no microbes competing for soil nutrients and less resources spent in attracting/repelling bacteria^32^). We found that, although the selected microbiomes’ positive effect on host fitness remained largely the same, selected microbiomes provided benefits to the host plants that were both stress-specific and non-specific. The precise mechanism behind the ‘non-specific’ benefit is unclear, but it could be partially due to loss of detrimental bacteria during successive selection cycles. The salt-specific benefit, on the other hand, is related to the positive selection of specific bacteria that either modify soil-nutrient availability or modulate plant-nutrient uptake.

Plants that received the sodium-selected microbiomes showed a ∼50% lower leaf-sodium concentration than those inoculated with microbiomes harvested from fallow-soil, which presumably explained the ∼140% higher total seed weight of these plants. The magnitude of this response is exceptional and probably driven by contribution of many taxa within the microbiome^33^. The precise mechanism underlying the action of each component is unknown, but bacteria-modulated salt tolerance has been linked to microbial production of phytohormones such as auxin^34,35^, cytokinin^36,37^, and gibberellin^38^. It has also been shown that some bacteria under salt stress produce 1-aminocyclopropane-1-carboxylate (AAC) deaminase, an enzyme that cleaves the ethylene precursor ACC, downregulating ethylene levels in the root and increasing plant growth^39^. High AAC deaminase activity has been observed in species of *Burkholderia*, *Pseudomonas*, *Rhizobium*, and *Mesorhizobium*^39^, the bacterial genera of several ASVs enriched in our sodium-selected microbiomes (Fig. 4e). In addition, some bacteria can produce exopolysaccharides that may help alleviate salt stress by binding with Na^+^ cations and reducing its availability for plant uptake^40^, whilst other bacteria may produce osmolytes^41^. Finally, bacteria can reduce the oxidative damage (production and accumulation of reactive oxygen species) of plants under salt stress by either producing antioxidant compounds^42^ or inducing the plant’s own antioxidant defense system^43^. This antioxidant response is especially interesting because it could benefit plants experiencing other abiotic stresses (including aluminium) and would explain, at least partially, the functional overlap we observed when selected microbiomes were subjected to an alternative stress in our cross-fostering experiment. In terms of aluminium stress, in addition to this non-specific response, we surmise that aluminium-selected microbiome lines also harboured bacterial taxa that produce exopolysaccharides and biofilms, which could potentially shield roots from high aluminium levels in the soil. This hypothesis is based on the phytotoxic mechanism of aluminium, which primarily affects cell division and elongation in the root apex^44,45^, and that increased mucilage production has been shown to alleviate aluminium toxicity in plants^46,47^. By excluding aluminium from root tips, either spatially or chemically (i.e., chelation by negatively charged compounds), these polymers could improve plant growth in a way that would not be evident by changes in leaf-aluminium concentration. Whether bacteria-produced exopolysaccharides display the same properties as root mucilage and provide the same effect is, however, unknown and warrants further study.

The design of our study allowed us to investigate how root-associated microbiome composition was altered after selection under distinct salt stresses. Our protocol implemented independent microbiome-selection “lines” (Supplementary Fig. S2), and selected microbiomes were not homogenized across different lines between growth cycles. A possibility is that throughout the microbiome-breeding process where different selection lines are not permitted to mix, multiple community modalities could emerge that would each confer tolerance to one stress, not necessarily due to the same bacterial components or through identical mechanisms. We did not observe this pattern, though: sodium- and aluminium-selected lines resulted in distinct communities clustered by the stress under which the microbiomes were selected (Fig. 4b), suggesting a common identity for these microbiomes across most lines that is rooted in both selection history and salt stress. The compositional differences between microbiomes of the two selection histories were characterized by differential abundance and extinction of microbial taxa, likely induced by a combination of direct response to the salt stress and indirect responses to plant cues, such as the “cry for help” response^48^.

Our approach for microbiome breeding is analogous to animal or crop breeding, which relies upon the inheritance of alleles that contribute to a desirable trait. However, unlike a genome, which is intrinsic to the plant, microbiomes are acquired from the environment, making microbiome breeding dependent on faithful transmission and assembly of the microbiome across generations. Because our microbiomes did not decay into monocultures throughout nine generations of selection, it appears that a complex community can be perpetuated with sufficient fidelity over generations with minimal invasion by environmentally-derived microbes. More importantly, our microbiomes remained intact with relatively minor perturbation in community composition even when challenged with an alternative salt stress (Fig. 4c). One likely explanation is that, as mentioned above, the breeding process eliminates bacterial taxa from the selection lines differently in response to different stresses, restricting the degree to which the microbiomes can shift to an alternative state. This might explain why cross-fostered microbiomes did not rescue plants to the same degree as microbiomes bred for the targeted stress (Fig. 2b). Together, these results suggest that the benefit conferred by artificially-selected microbiome is at least partially explained by its community composition. An open question that remains to be addressed is to what degree within-strain genomic evolution imparts host protection against the imposed stresses.

In addition to changes in microbial community composition, we also observed significant changes in co-occurrence patterns among bacterial species with our selection protocol (Fig. 5). The increase in frequency and strength of correlations among microbiome components with selection history and salt stress could point towards an increase in biotic interactions among bacteria, a phenomenon that might help stabilize these communities and explain their compositional robustness in the face of perturbation (i.e., switch to alternative salt stress during cross-fostering), as well as their high transmission fidelity and heritability across selections rounds. Because networks were reconstructed using a correlation-based approach based on taxa occurrence and abundance data, conclusions from network features regarding mechanisms underlying transmission fidelity need to be inferred with caution. From an ecological standpoint, species co-occurrence patterns could be driven by stochastic processes, such as dispersal limitation and drift^49,50^, or by deterministic processes, such as environmental filtering and biotic interactions^51,52^. Our selection protocol aimed to minimize the impact of stochastic processes through inoculation of plants with a large volume of transferred microbiomes, thus maximizing priority effects when these establish around the roots, the nearly exclusion of bacterial dispersion across communities, and the slow ramping of imposed stresses between microbiome generations^17^, which suggests that the observed differences in network structure and topology were due to deterministic processes. Disentangling the relative importance of environmental filtering and biotic interactions during the selection step is difficult because both processes can play a role in shaping communities^53^, but the results of our network analyses do suggest that biotic interactions are involved: differences between the selected microbiomes’ networks under their corresponding and alternative salt stress can only derive from environmental filtering, which means that biotic interactions likely drive at least part of the difference in edge density between networks when selection history matched salt stress (i.e., highest complexity) and the null networks. The increase in clustering coefficient and, more importantly, natural connectivity also points towards an increase in biotic interactions. It is highly unlikely that such a discrepancy in global clustering coefficient among networks of the same size (i.e., same number of nodes) is a mathematical artifact given its strong correlation with selection history, even considering the compositional nature of the data. Because the clustering coefficient is related to the ‘cliquishness’ of nodes in a graph^54^ – that is, their grouping in closely connected subsets – an increase in this clustering metric suggests that more bacterial species are interacting, forming ‘cliques’ that are likely functionally determined. As for natural connectivity, this metric has been proposed as a simple and effective measurement of network robustness (i.e., the ability of a network to maintain its connectivity when a fraction of its vertices is lost) because it changes strictly monotonically with the addition or deletion of edges^25^. This means that any change in natural connectivity is rooted in a change of the inherent structure of the network and not due to spurious correlations from the compositional data. The increase in natural connectivity - and network robustness - that we observed when salt stress matched the selection history is, therefore, at least partially a consequence of an increase in real interactions among bacteria species in those communities. Moreover, exploring co-occurrence patterns as an additional dimension of the microbial communities allowed us to identify candidate bacteria (or groups of bacteria) that, despite not being necessarily the most abundant, could disproportionally influence community structure and its functioning. This is interesting from a hypothesis-ranking perspective, providing a list of microbes that are more likely to be causal to the observed phenotypes or have a higher centrality, thus being potentially useful in a synthetic microbiome framework.

In summary, our findings establish host-mediated artificial selection as an efficient and robust method for breeding microbiomes with specific functions, particularly when the selection scheme considers aspects relative to plant control^55^ over microbiome and the heritability of microbial effects. These aspects, combined with a sufficiently high fidelity in microbiome transmission between plants, ties the fitness interests of microbes to those of their host^55^, enabling host-mediated selection. With our protocol, we were able to select microbiomes that increased plant tolerance to sodium and aluminium stress, two major agricultural problems that are expected to worsen as climate changes. To deal with such stresses, plant breeders have been working on stress-tolerant genotypes, but phenotypic selection and breeding approaches, even modern ones involving marker-assisted and genomic selection, are slow. With our microbiome breeding protocol, we were able to achieve sizeable results (up to ∼200% increase in seed output) surprisingly fast, within 1-3 microbiome-selection cycles. It is possible that the microbiome-encoded effects selected in one species might be, at least partially, transferable to other species, which would broaden the agricultural applications of microbiome breeding.

## Methods

### Plant material and microbiome-selection scheme

Figure 1 illustrates the microbiome-selection scheme. *Brachypodium distachyon* (Bd3-1) seeds from a non-evolving seedstock were used throughout the nine selection cycles and in the final 2×2 partial cross-fostering experiment at Generation 9. During the selection cycles (i.e., the microbiome breeding phase), five selection lines were established per treatment (aluminium and sodium stress) and fallow-soil controls, each line comprised of eight individual plants (or root-free soil) for a total of 40 replicates per treatment per generation. The plants were inoculated with a diverse microbiome composed of root-associated bacteria collected from greenhouse-grown plants of *B. distachyon* and five local grass species^17^. Microbiomes were allowed to mature and interact with plants for ∼28 days, then plants within each selection line were visually phenotyped by size, with the two largest plants selected for microbiome harvest; their roots were submerged in extraction buffer and gently shaken until most material adhering to the root was released into solution, which was then filtered to remove soil particles and larger-sized organisms (e.g., fungi, algae) that could otherwise co-propagate with the bacterial fraction, the target of our microbiome-selection protocol. The solution obtained from the two largest plants within each selection line were pooled to create the microbiome used to inoculate plants hosting the subsequent microbiome-generation, established from the same seedstock, and grown in the same media (Profile® porous ceramic rooting media), but with a higher concentration of either aluminium or sodium sulfate (see Mueller *et al*, 2021^17^ for concentrations used in the experiment). This ramping of salt stress was incorporated for two reasons: 1. to reduce the effect of drift on microbiome evolution; and 2. to impose a significant, yet moderately increasing, level of stress throughout the selection cycles, as overstressed plants may not “feel” the impact of beneficial microbes, whilst understressed plants may not need the help of beneficial microbes. The microbiome breeding phase lasted for nine generations (i.e., the selection cycles illustrated in Fig. 1 and described in detail in Mueller et al^17^). After nine generations, we tested the effect of the resulting microbiomes in plants experiencing sodium and aluminium salt stress using a 2×2 partial cross-fostering experiment (Fig. 2a) where seedlings were subject to one of the following treatments: 1) aluminium-selected microbiomes that were 2µm-filtered to propagate the bacterial microbiome fraction; 2) sodium-selected microbiomes that were 2µm-filtered to propagate the bacterial microbiome fraction; 3) solutes from aluminium- or sodium-selected microbiomes, where selected microbiomes were double-filtered (2µm and then 0.2μm) to remove live bacterial cells, so any effect of this filtrate on plant growth would be the result of only root exudates or viruses that may passed through the 0.2μm filter; 4) fallow-soil microbiomes, where bacterial microbiomes were harvested from root-free fallow soil subject to the same ramping of the imposed stresses through nine generations; and, 5) no bacterial inoculum.

### DNA extraction, 16S rRNA amplicon sequencing, and ASV identification

Genomic DNA was extracted from filtered washes of root systems using the DNeasy PowerSoil Pro Kit (QIAGEN, Germany). DNA libraries for sequencing of microbial 16S rRNA V4 and V4-V5 gene regions were prepared with a PerkinElmer Sciclone NGS robotic liquid handling system using 30ng sample input, custom-designed target primers (515F/806R and 515F-Y/926R for the V4 and V4-V5 gene regions, respectively) with incorporated Illumina sequencing adapters with unique indexes (IDT), chloroplast and mitochondrial PNA blocking oligos for 16S endophyte samples (PNA Bio), and 5 PRIME HotMasterMix® Taq DNA Polymerase (Quantabio). Libraries were then pooled together, and the pool was quantified on a Roche LightCycler® 480 real-time PCR Instrument II using KAPA Biosystem’s NGS qPCR-based library quantification kit (Roche). The library pool was sequenced at the DOE Joint Genome Institute (Walnut Creek, https://jgi.doe.gov/) on an Illumina MiSeq Sequencing Platform using the MiSeq Reagent Kit v3 (600-cycles), following a 2×300 indexed-run recipe. The reads were trimmed to remove primer and adapter sequences using Cutadapt^56^. Sequences were truncated to 240 bp, filtered for a maximum of 4 expected errors, denoised, and clustered into ASVs using Dada2^57^. The resulting table was filtered to remove chimeric ASVs, and taxonomies of the sequences were assigned against the Silva database version 13.2^58^.

### Microbiome community composition analyses

The Generation 9 ASV table was converted to relative abundances by dividing raw counts by the library sequencing depth. ASVs detected in less than 10% of the samples were excluded from further analysis. Principal coordinates analyses were done using the *capscale* function, Shannon diversity was calculated using the *diversity* function, and beta diversity within sample types was calculated using the *betadisper* function, all from the vegan^59^ package. Differential abundance of the bacterial taxa between selection histories was estimated using a linear modelling approach where the log2-transformed relative abundance of each microbe was modelled as a function of selection history. Only ASVs present in ≥25% of the samples were retained for the differential abundance analysis.

### Microbial co-occurrence network construction

Bacterial co-occurrence networks were estimated for each selection history (sodium, aluminium, and null) and salt stress combination. The networks were restricted to a maximum size of 50 nodes (i.e., the 50 most abundant bacterial ASVs) as this represented 98.1-99.9% of reads in each network whilst still allowing them to be interpretable. Using SparCC^2^, we estimated the approximated linear Pearson correlations between log-ratio transformed ASV fractions, then constructed the adjacency matrix using Student’s t-test as a sparsification method. The resulting networks were analysed for number of components, clustering coefficient, modularity, edge density, positive edge percentage, and natural connectivity using NetCoMi^20^ (Table S1). In addition to these metrics, we also calculated average dissimilarity and average path length for the largest connected component (LCC) of each network (Table S2).

### Within-module degree and participation coefficient

To classify nodes according to their topological properties in each network, we calculated their patterns of connectivity both within (within-module degree, *z*) and across modules (participation coefficient, *P*). *z_i_* measures how connected node *i* is to other nodes in its own module, such that:

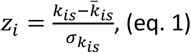

where *k_is_* is the number of edges from node *i* to all other nodes in module *s*, 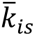 is the average of *k* across all nodes in *s*, and

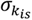 is the standard deviation of *k* in *s*. As for the participation coefficient *P*, it measures how distributed the edges of node *i* are among different modules, and was defined as:

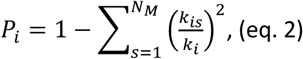

where *k_is_* is the number of edges of node *i* to other nodes in module *s*, and *k_i_* is the total degree of node *i*. Each node was then heuristically assigned a role depending on their placement in the *z*-*P* parameter space^61^ (Fig. 3c), with nodes with a *z*≥1.5 being defined as hubs and those with a *z*<1.5 as non-hubs. Both hub and non-hub nodes were more finely characterized by their participation coefficient: hub nodes with *P*≥0.62 were classified as network hubs; hub nodes with *P*<0.62 were classified as module hubs; non-hub nodes with *P*≥0.62 were classified as connectors; and non-hub nodes with *P*<0.62 were classified as peripherals.

### Leaf-tissue nutrient analysis

After collecting the microbiome samples of plants at generation 9, we harvested the aboveground tissue of each individual plant and oven-dried it at 60°C for >72h. We then ground the leaf material (SPEX SamplePrep Geno-Grinder 2010) using plastic vials and yttria-stabilized zirconium beads and analysed the tissue element concentration through inductively coupled plasma mass spectrometry (ICP-MS; Perkin-Elmer SCIEX Elan 6000 DRC-e mass spectrometer) connected to a PFA microflow nebulizer (Elemental Scientific) and an Apex HF desolvator (Elemental Scientific). In total, 400 leaf samples were analysed for aluminium and sodium (Al and Na, respectively; Fig. 3a), magnesium, phosphorus, potassium, and calcium (Mg, P, K, and Ca, respectively; Supplementary Fig. S3), boron, iron, manganese, nickel, copper, zinc, and molybdenum (B, Fe, Mn, Ni, Cu, Zn, and Mo, respectively; Supplementary Fig. S4), cobalt, arsenic, selenium, rubidium, strontium, and cadmium (Co, As, Se, Rb, Sr, and Cd, respectively; Supplementary Fig. S5).

### Statistical analyses

Differences in total seed weight (Fig. 2) and leaf-tissue nutrient concentration (Figs 3, S3-S5) were analysed using generalized linear mixed-effect models^62^ (GLMMs). Models were fitted to the data considering the parameter to be analysed as function of the bacterial treatment, with selection line as a random effect (see Tables S3-S7 for details). The significant models were selected using Akaike Information Criterion corrected for finite sample sizes (AICc), then screened for homoscedasticity and normality of the residuals, with appropriate variance structures applied whenever necessary^63^. Differences between bacterial treatment were determined posthoc through Šídák multiple-comparison tests. The linear mixed-effect models were created and analysed with the lme4^64^ package. The correlations were performed using mean values of total seed weight (fitness) and leaf-nutrient concentrations for each selection line, and thus intrinsically estimated with error. Because of this, we employed a model-II simple linear regression analysis using ordinary-least squares with the lmodel2 package. All statistical analyses were performed in the R Environment^65^.

## Resource availability

The host-mediated artificial-selection protocol has been published in detail (https://doi.org/10.1128/mSystems.01125-21). The raw sequencing data - including 16S rRNA sequence - generated in this study have been deposited at the NCBI database under BioProject accession no. XXXXXXXXXXX (https://www.ncbi.nlm.nih.gov/bioproject/?term=XXXXXXXXXXX). Source data can be accessed at Dryad (DOI: 10.5061/dryad.h44j0zpwg), and code used to perform the analyses are available on Zenodo (https://doi.org/10.5281/zenodo.14232830).

### Acknowledgements

We thank Mark Seniw for designing Figure 1. The research was funded by Department of Energy award DE-SC0021126, Joint Genome Institute CSP award #1573, and the Stengl Lost-Pines Endowment from The University of Texas at Austin. The work (Proposal: Award DOI 10.46936/10.25585/60000712) conducted by the U.S. Department of Energy Joint Genome Institute (https://ror.org/04xm1d337), a DOE User Facility, is supported by the Office of Science of the U.S. Department of Energy operated under Contract No. DE-AC02-05CH11231.

## Author Contributions

Conceptualization, U.G.M.; Investigation, C.G.P., J.A.E., A.K., A.C., and U.G.M.; Writing – Original Draft, C.G.P., J.A.E., and U.G.M.; Writing – Review & Editing, C.G.P., J.A.E., A.K., T.G.R., J.P.V., D.L.D.M., T.E.J., and U.G.M.; Funding Acquisition, T.E.J. and U.G.M.; Resources, V.B., E.S., T.G.R., S.T., J.P.V., D.L.D.M., T.E.J., and U.G.M.; Supervision, T.E.J. and U.G.M.

## Supplementary data

**Fig S1.** Shannon diversity of aluminium- and sodium-selected microbiomes under both aluminium and sodium stress.

**Fig S2.** Principal coordinate analyses (PCoA) of selected microbiome lines showing clustering based on selection history.

**Fig S3.** *z*-*P* plots for each microbial co-occurrence network of root-associated bacteria after 9 generations of artificial selection.

**Fig S4.** Leaf macronutrient (Mg, P, K, Ca) concentration of plants given different inocula at our cross-fostering experiment.

**Fig S5.** Leaf micronutrient (B, Fe, Mn, Ni, Cu, Zn, Mo) concentration of plants given different inocula at our cross-fostering experiment.

**Fig S6.** Leaf trace-element (Co, As, Se, Rb, Sr, Cd) concentration of plants different inocula at our cross-fostering experiment.

**Table S1.** Largest connected component (LCC) parameters of *B. distachyon* root-associated bacteria co-occurrence networks.

**Table S2.** Linear mixed-effect models for the analysis of total seed weight (Fig. 2b) of plants in our cross-fostering experiment.

**Table S3.** Linear mixed-effect models for the analysis of leaf Al and Na concentration (Fig. 3) of plants in our cross-fostering experiment.

**Table S4.** Linear mixed-effect models for the analysis of leaf macronutrient concentration (Fig. S4) of plants in our cross-fostering experiment.

**Table S5.** Linear mixed-effect models for the analysis of leaf micronutrient concentration (Fig. S5) of plants in our cross-fostering experiment.

**Table S6.** Linear mixed-effect models for the analysis of leaf trace-element concentration (Fig. S6) of plants in our cross-fostering experiment.

