## Supplemental Information for "Breeding of microbiomes conferring salt tolerance to plants"

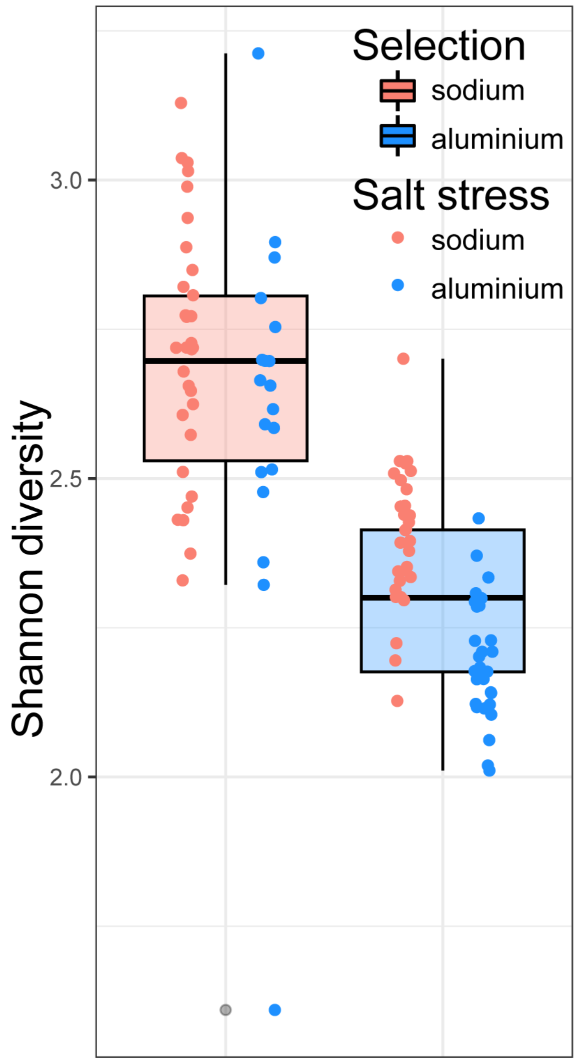


**Figure S1. Microbiome breeding through host-mediated artificial selection impacts alpha-diversity of microbiomes differently depending on the imposed salt stress.** Shannon diversity estimates for root-associated microbial communities of plants given sodium- and aluminium-selected microbiomes under sodium and aluminium salt stresses. Selection under sodium stress resulted in more diverse microbiomes than those selected under aluminium stress.

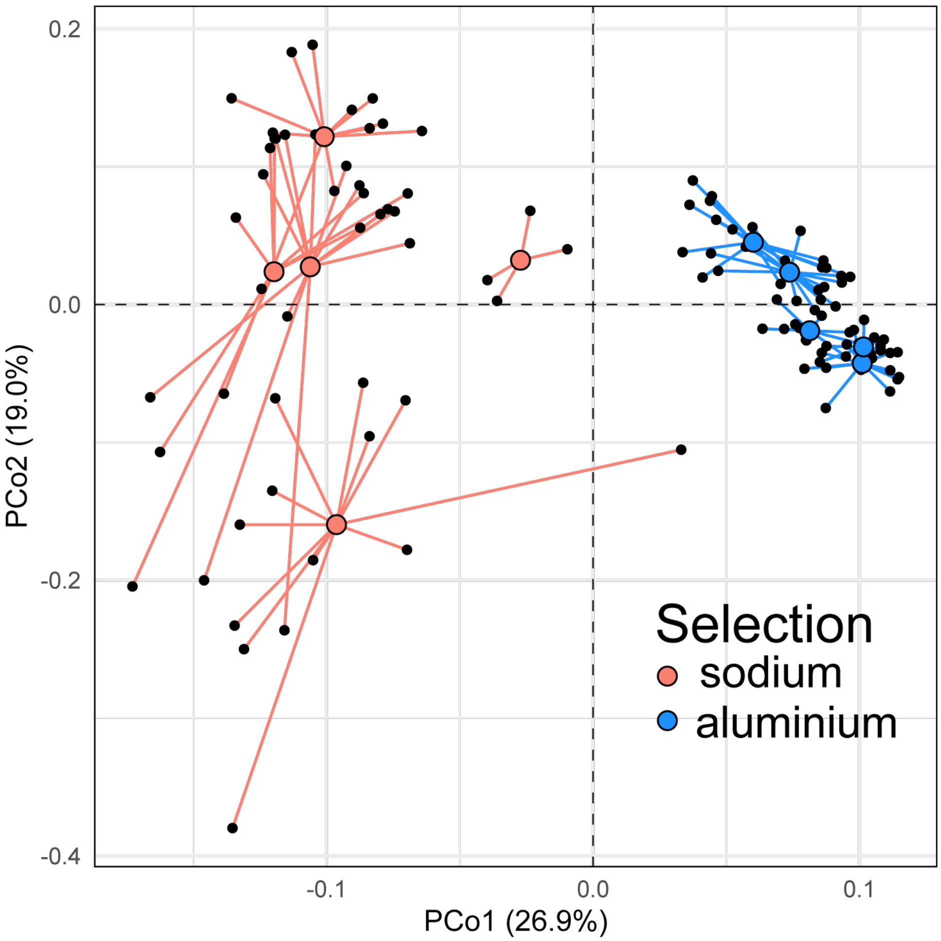


**Figure S2. Different microbiome selection lines show similar composition within selection history.** Principal coordinate analysis (PCoA) based on Bray-Curtis dissimilarity showing distinct clusters of root-associated microbial communities based on selection history in our cross-fostering experiment (Fig. 4b). The proportion of variance explained by each PCoA is denoted in the corresponding axis labels. The centroids for each selection line are larger, coloured dots (red for sodium-selected lines, blue for aluminium-selected lines) connected to individual samples (in black).

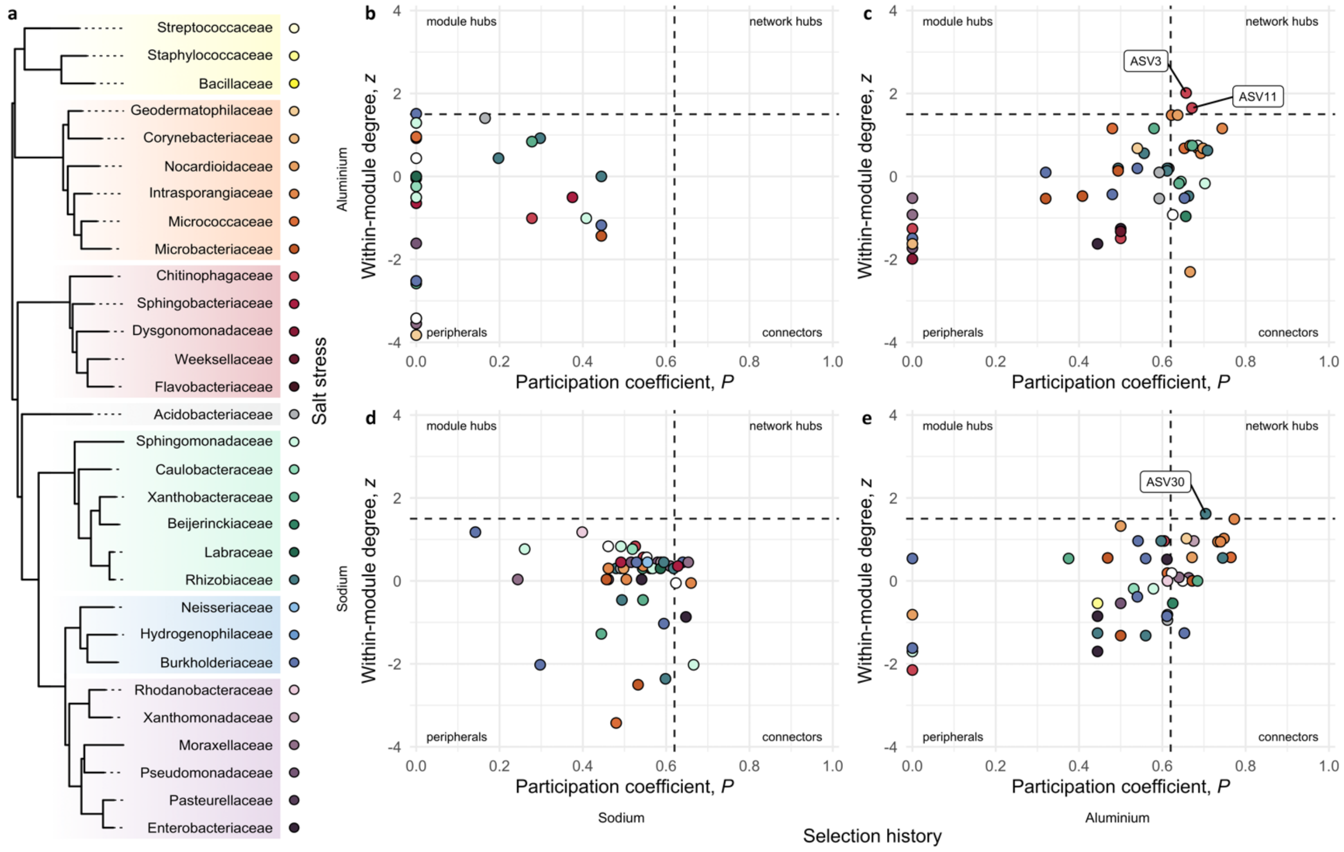


**Figure S3. Microbial co-occurrence networks for root-associated bacteria after 9 generations of artificial selection. (a)** Phylogenetic relationship among bacterial families of ASVs identified in our network analyses, as extracted from the maximum-likelihood estimated molecular phylogeny including Bacteria, Archaea and Eukarya[64]; each family is represented by a unique colour, with different hues depicting classes: yellow for Bacilli, orange for Actinobacteria, red for Bacteroidia, grey for Acidobacteriia, green for α-Proteobacteria, blue for β -Proteobacteria, purple for γ-Proteobacteria. **(b-e)** *z-P* plot exhibiting patterns of within- and across-module connectivity of ASVs from the microbial co-occurrence networks inferred for plants given sodium- and aluminium-selected microbiomes under sodium and aluminium stress, with the identified network hubs highlighted.

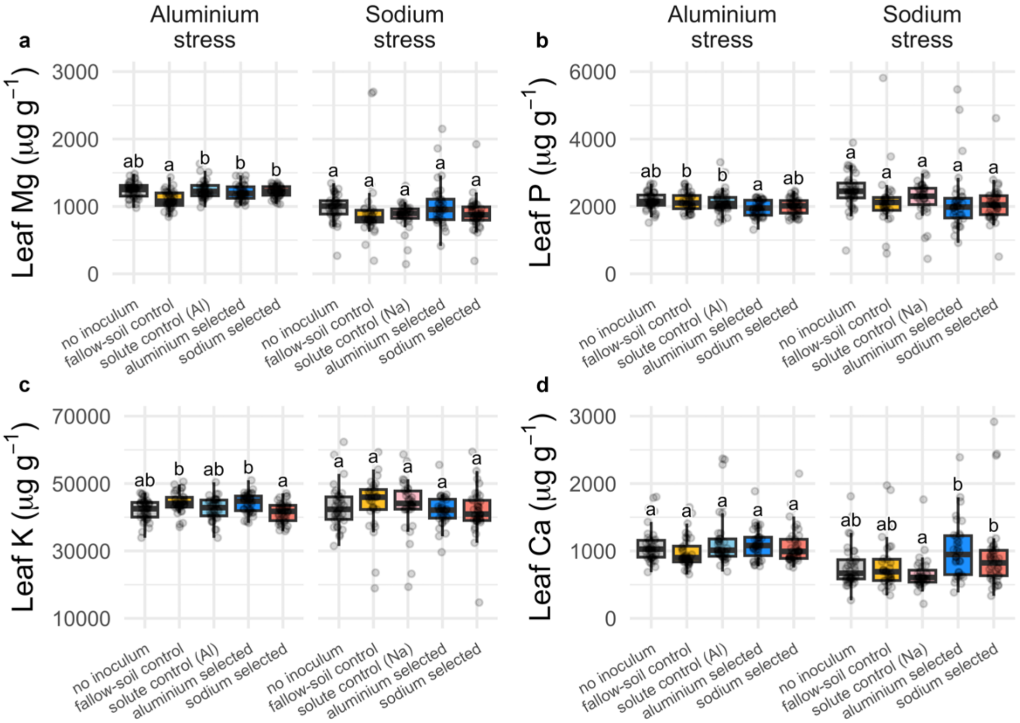


**Figure S4. Leaf macronutrient concentration of plants given different inocula in our cross-fostering experiment. (a)** Magnesium (Mg); **(b)** phosphorus (P); **(c)** potassium (K); and **(d)** calcium (Ca). Boxplots with medians, 25$^{th}$, and 75$^{th}$ percentiles. Whiskers extend to 1.5 times the interquartile range and data points presented beyond whiskers represent outliers. Different letters indicate significant differences between bacterial treatments within individual salt stresses based on Šídák posthoc tests.

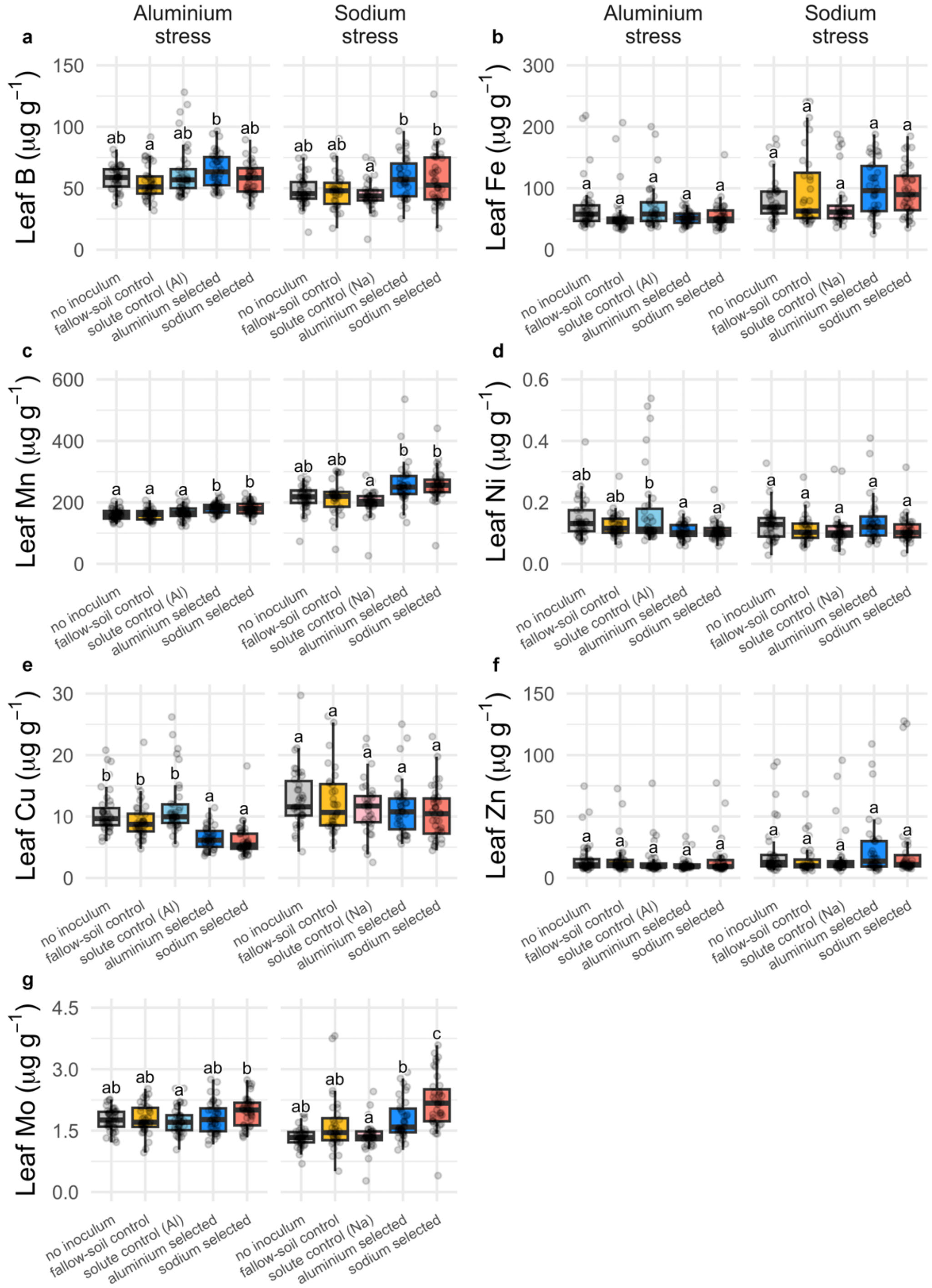


**Figure S5. Leaf micronutrient concentration of plants given different inocula in our cross-fostering experiment. (a)** Boron (B); **(b)** iron (Fe); **(c)** manganese (Mn); **(d)** nickel (Ni); **(e)** copper (Cu); **(f)** zinc (Zn); and **(g)** molybdenum (Mo). Boxplots with medians, 25$^{th}$, and 75$^{th}$ percentiles. Whiskers extend to 1.5 times the interquartile range and data points presented beyond whiskers represent outliers. Different letters indicate significant differences between bacterial treatments within individual salt stresses based on Šídák posthoc tests.

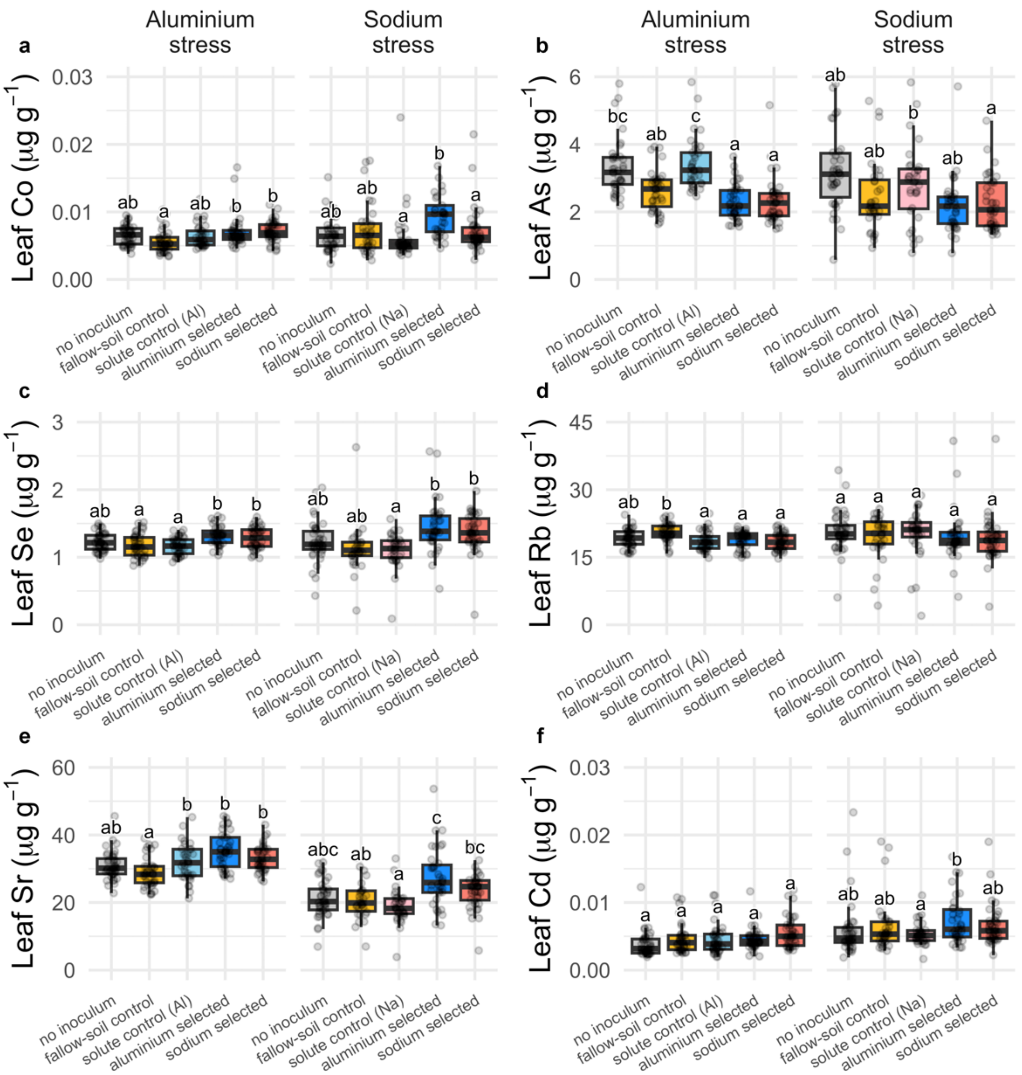


**Figure S6. Leaf trace-element concentration of plants different inocula in our cross-fostering experiment. (a)** Cobalt (Co); **(b)** arsenic (As); **(c)** selenium (Se); **(d)** rubidium (Rb); **(e)** strontium (Sr); and **(f)** cadmium (Cd). Boxplots with medians, 25th, and 75th percentiles. Whiskers extend to 1.5 times the interquartile range and data points presented beyond whiskers represent outliers. Different letters indicate significant differences between bacterial treatments within individual salt stresses based on Šídák posthoc tests.

**Table S1.** Network parameters for the largest connected component (LCC) of *B. distachyon* (Bd3-1) associated root microbiome at two salt stresses (aluminium and sodium sulphate) after 9 rounds of artificial selection for sodium and aluminium resistance, as well as the control (null), which has not been selected. The "sodium stress, null selection history" network was empty, thus the absence of results.

| Salt stress | Selection history | LCC Size | Clustering coefficient | Modularity | Positive edge percentage | Edge density | Natural connectivity | Average dissimilarity | Average path length |
|---|---|---|---|---|---|---|---|---|---|
| **aluminium** | null | 0.44 | 0.00 | 0.19 | 93.33 | 0.13 | 0.16 | 0.88 | ∞ |
| | sodium | 0.48 | 0.72 | 0.44 | 91.30 | 0.25 | 0.13 | 0.86 | 1.46 |
| | aluminium | 0.96 | 0.45 | 0.21 | 50.00 | 0.22 | 0.11 | 0.90 | ∞ |
| **sodium** | null | - | - | - | - | - | - | - | - |
| | sodium | 1 | 0.74 | 0.13 | 55.20 | 0.54 | 0.19 | 0.81 | 1.06 |
| | aluminium | 0.98 | 0.49 | 0.18 | 43.52 | 0.22 | 0.08 | 0.92 | 1.32 |

**Table S2. Linear mixed-effect models for the analysis of total seed weight** (Fig. 2b) **of plants under each salt stress in our cross-fostering experiment.** Data analysed, model identification and rationale, number of parameters (*K*), log likelihood (log L), Akaike Information Criterion corrected for finite sample sizes (AICc), difference between selected and null model`s AICc (ΔAICc), and *p*-value. The significantly different values (ANOVA) are indicated: ***, significant ($p < 0.05$); and ns, non-significant. The identification after model ID (e.g., _*vs1*) refers to the variance structure applied to the model in order to satisfy the prerequisites of heteroscedasticity and normality of residues. Using the *nlme* package in R, we used: _*vs1*, varIdent(~1|Treat); _*vs2*, varPower(); _*vs3*, varPower(~fitted(.)|Treat); _*vs4*, varConstPower(); _*vs5*, varComb(varIdent(~1|Treat), varPower()); _*vs6*, varExp(); and _*vs7*, varExp(~fitted(.)|Treat).

| Data | Model ID | Model | *K* | Log L | AICc | ΔAICc | *p* |
|---|---|---|---|---|---|---|---|
| **Seed weight** (Al stress; fig. 2b) | mm1_vs6 | Seed weight ~ Treat \| random = Selection line | 8 | -1154.21 | 2325.20 | -45.17 | <0.001 *** |
| | null | Seed weight ~ 1 \| random = Selection line | 3 | -1182.12 | 2370.37 | - | - |
| **Seed weight** (Na stress; fig. 2b) | mm1_vs6 | Seed weight ~ Treat \| random = Selection line | 8 | -1148.71 | 2314.23 | -51.01 | <0.001 *** |
| | null | Seed weight ~ 1 \| random = Selection line | 3 | -1179.55 | 2365.24 | - | - |

**Table S3. Linear mixed-effect models for the analysis of leaf aluminium and sodium concentration** (Al and Na, respectively; Fig. 3a) **of plants under each salt stress in our cross-fostering experiment.** Data analysed, model identification and rationale, number of parameters (*K*), log likelihood (log L), Akaike Information Criterion corrected for finite sample sizes (AICc), difference between selected and null model`s AICc (ΔAICc), and *p*-value. The significantly different values (ANOVA) are indicated: $^{***}$, significant ($p < 0.05$); and $^{ns}$, non-significant. The identification after model ID (e.g., *_vs1*) refers to the variance structure applied to the model in order to satisfy the prerequisites of heteroscedasticity and normality of residues. Using the *nlme* package in R, we used: *_vs1*, varIdent(~1|Treat); *_vs2*, varPower(); *_vs3*, varPower(~fitted(.)|Treat); *_vs4*, varConstPower(); *_vs5*, varComb(varIdent(~1|Treat), varPower()); *_vs6*, varExp(); and *_vs7*, varExp(~fitted(.)|Treat).

| Data | Model ID | Model | *K* | Log L | AICc | ΔAICc | *p* |
|---|---|---|---|---|---|---|---|
| **Leaf Al** (Al stress; fig. 3a) | mm1_vs2 | Leaf Al ~ Treat \| random = Selection line | 8 | -215.18 | 447.17 | -15.76 | <0.001 $^{***}$ |
| | null | Leaf Al ~ 1 \| random = Selection line | 3 | -228.40 | 462.93 | - | - |
| **Leaf Al** (Na stress; fig. 3a) | mm1_vs1 | Leaf Al ~ Treat \| random = Selection line | 11 | -295.07 | 613.73 | -17.34 | <0.001 $^{***}$ |
| | null | Leaf Al ~ 1 \| random = Selection line | 3 | -312.47 | 631.07 | - | - |
| **Leaf Na** (Al stress; fig. 3a) | mm1_vs1 | Leaf Na ~ Treat \| random = Selection line | 11 | -495.22 | 1013.92 | -90.79 | <0.001 $^{***}$ |
| | null | Leaf Na ~ 1 \| random = Selection line | 3 | -549.29 | 1104.71 | - | - |
| **Leaf Na** (Na stress; fig. 3a) | mm1_vs6 | Leaf Na ~ Treat \| random = Selection line | 8 | -1206.74 | 2430.35 | -26.9 | 0.017 $^{***}$ |
| | null | Leaf Na ~ 1 \| random = Selection line | 3 | -1225.55 | 2457.25 | - | - |

**Table S4. Linear mixed-effect models for the analysis of leaf macronutrient** (Mg, P, K, and Ca; Fig. S4) **concentration of plants under each salt stress in our cross-fostering experiment.** Data analysed, model identification and rationale, number of parameters (*K*), log likelihood (log L), Akaike Information Criterion corrected for finite sample sizes (AICc), difference between selected and null model`s AICc (ΔAICc), and *p*-value. The significantly different values (ANOVA) are indicated: $^{***}$, significant ($p < 0.05$); and $^{ns}$, non-significant. The identification after model ID (e.g., *_vs1*) refers to the variance structure applied to the model in order to satisfy the prerequisites of heteroscedasticity and normality of residues. Using the *nlme* package in R, we used: *_vs1*, varIdent(~1 |Treat); *_vs2*, varPower(); *_vs3*, varPower(~fitted(.)|Treat); *_vs4*, varConstPower(); *_vs5*, varComb(varIdent(~1|Treat), varPower()); *_vs6*, varExp(); and *_vs7*, varExp(~fitted(.)|Treat).

| Data | Model ID | Model | *K* | Log L | AICc | ΔAICc | *p* |
|---|---|---|---|---|---|---|---|
| **Leaf Mg** (Al stress; fig. S1) | mm1 | Leaf Mg ~ Treat \| random = Selection line | 7 | -1162.44 | 2339.50 | -24.51 | <0.001 $^{***}$ |
| | null | Leaf Mg ~ 1 \| random = Selection line | 3 | -1178.94 | 2364.01 | - | - |
| **Leaf Mg** (Na stress; fig. S1) | mm1_vs1 | Leaf Mg ~ Treat \| random = Selection line | 11 | -1241.49 | 2506.58 | -32.85 | <0.001 $^{***}$ |
| | null | Leaf Mg ~ 1 \| random = Selection line | 3 | -1266.64 | 2539.43 | - | - |
| **Leaf P** (Al stress; fig. S1) | mm1 | Leaf P ~ Treat \| random = Selection line | 7 | -1335.95 | 2686.52 | -9.65 | 0.0012 $^{***}$ |
| | null | Leaf P ~ 1 \| random = Selection line | 3 | -1345.02 | 2696.17 | - | - |
| **Leaf P** (Na stress; fig. S1) | mm1_vs1 | Leaf P ~ Treat \| random = Selection line | 11 | -1443.64 | 2910.87 | -26.42 | <0.001 $^{***}$ |
| | null | Leaf P ~ 1 \| random = Selection line | 3 | -1465.58 | 2937.29 | - | - |
| **Leaf K** (Al stress; fig. S1) | mm1 | Leaf K ~ Treat \| random = Selection line | 7 | -1778.15 | 3570.92 | -10.99 | <0.001 $^{***}$ |
| | null | Leaf K ~ 1 \| random = Selection line | 3 | -1787.89 | 3581.91 | - | - |
| **Leaf K** (Na stress; fig. S1) | mm1 | Leaf K ~ Treat \| random = Selection line | 7 | -1791.40 | 3597.49 | 7.97 | 0.9656 $^{ns}$ |
| | null | Leaf K ~ 1 \| random = Selection line | 3 | -1791.69 | 3589.52 | - | - |
| **Leaf Ca** (Al stress; fig. S1) | mm1_vs1 | Leaf Ca ~ Treat \| random = Selection line | 11 | -1328.25 | 2679.99 | -9.52 | <0.001 $^{***}$ |
| | null | Leaf Ca ~ 1 \| random = Selection line | 3 | -1341.69 | 2689.51 | - | - |
| **Leaf Ca** (Na stress; fig. S1) | mm1_vs2 | Leaf Ca ~ Treat \| random = Selection line | 8 | -1286.67 | 2590.21 | -20.78 | <0.001 $^{***}$ |
| | null | Leaf Ca ~ 1 \| random = Selection line | 3 | -1302.43 | 2610.99 | - | - |

**Table S5. Linear mixed-effect models for the analysis of leaf micronutrient** (B, Fe, Mn, Ni, Cu, Zn, and Mo; Fig. S5) **concentration of plants under each salt stress in our cross-fostering experiment.** Data analysed, model identification and rationale, number of parameters (*K*), log likelihood (log L), Akaike Information Criterion corrected for finite sample sizes (AICc), difference between selected and null model`s AICc (ΔAICc), and *p*-value. The significantly different values (ANOVA) are indicated: $^{***}$, significant ($p < 0.05$); and $^{ns}$, non-significant. The identification after model ID (e.g., _*vs1*) refers to the variance structure applied to the model in order to satisfy the prerequisites of heteroscedasticity and normality of residues. Using the *nlme* package in R, we used: _*vs1*, varIdent(~1|Treat); _*vs2*, varPower(); _*vs3*, varPower(~fitted(.)|Treat); _*vs4*, varConstPower(); _*vs5*, varComb(varIdent(~1|Treat), varPower()); _*vs6*, varExp(); and _*vs7*, varExp(~fitted(.)|Treat).

| Data | Model ID | Model | *K* | Log L | AICc | ΔAICc | *p* |
|---|---|---|---|---|---|---|---|
| **Leaf B** (Al stress; fig. S2) | mm1 | Leaf B ~ Treat \| random = Selection line | 7 | -789.42 | 1593.45 | -1.65 | 0.0383 $^{***}$ |
| | null | Leaf B ~ 1 \| random = Selection line | 3 | -794.48 | 1595.10 | - | - |
| **Leaf B** (Na stress; fig. S2) | mm1 | Leaf B ~ Treat \| random = Selection line | 7 | -733.04 | 1480.75 | -10.09 | <0.001 $^{***}$ |
| | null | Leaf B ~ 1 \| random = Selection line | 3 | -742.35 | 1490.84 | - | - |
| **Leaf Fe** (Al stress; fig. S2) | mm1_vs1 | Leaf Fe ~ Treat \| random = Selection line | 11 | -874.56 | 1772.64 | -54.45 | <0.001 $^{***}$ |
| | null | Leaf Fe ~ 1 \| random = Selection line | 3 | -910.48 | 1827.09 | - | - |
| **Leaf Fe** (Na stress; fig. S2) | mm1 | Leaf Fe ~ Treat \| random = Selection line | 7 | -914.21 | 1843.08 | 0.47 | 0.0894 $^{ns}$ |
| | null | Leaf Fe ~ 1 \| random = Selection line | 3 | -918.24 | 1842.61 | - | - |
| **Leaf Mn** (Al stress; fig. S2) | mm1 | Leaf Mn ~ Treat \| random = Selection line | 7 | -829.73 | 1674.08 | -30.84 | <0.001 $^{***}$ |
| | null | Leaf Mn ~ 1 \| random = Selection line | 3 | -849.40 | 1704.92 | - | - |
| **Leaf Mn** (Na stress; fig. S2) | mm1_vs1 | Leaf Mn ~ Treat \| random = Selection line | 11 | -975.82 | 1975.24 | -63.67 | <0.001 $^{***}$ |
| | null | Leaf Mn ~ 1 \| random = Selection line | 3 | -1016.39 | 2038.91 | - | - |
| **Leaf Ni** (Al stress; fig. S2) | mm1_vs6 | Leaf Ni ~ Treat \| random = Selection line | 8 | 297.39 | -577.97 | -174.05 | <0.001 $^{***}$ |
| | null | Leaf Ni ~ 1 \| random = Selection line | 3 | 205.03 | -403.92 | - | - |
| **Leaf Ni** (Na stress; fig. S2) | mm1_vs6 | Leaf Ni ~ Treat \| random = Selection line | 8 | 260.91 | -504.96 | -6.53 | 0.004 $^{***}$ |
| | null | Leaf Ni ~ 1 \| random = Selection line | 3 | 252.29 | -498.43 | - | - |
| **Leaf Cu** (Al stress; fig. S2) | mm1_vs1 | Leaf Cu ~ Treat \| random = Selection line | 11 | -472.43 | 968.35 | -89.33 | <0.001 $^{***}$ |
| | null | Leaf Cu ~ 1 \| random = Selection line | 3 | -525.77 | 1057.68 | - | - |
| **Leaf Cu** (Na stress; fig. S2) | mm1 | Leaf Cu ~ Treat \| random = Selection line | 7 | -541.02 | 1096.69 | 2.23 | 0.1782 $^{ns}$ |
| | null | Leaf Cu ~ 1 \| random = Selection line | 3 | -544.16 | 1094.46 | - | - |
| **Leaf Zn** (Al stress; fig. S2) | mm1_vs2 | Leaf Zn ~ Treat \| random = Selection line | 8 | -735.96 | 1488.71 | -19.82 | <0.001 $^{***}$ |
| | null | Leaf Zn ~ 1 \| random = Selection line | 3 | -751.20 | 1508.53 | - | - |
| **Leaf Zn** (Na stress; fig. S2) | mm1_vs1 | Leaf Zn ~ Treat \| random = Selection line | 11 | -799.53 | 1622.65 | -9.4 | <0.001 $^{***}$ |
| | null | Leaf Zn ~ 1 \| random = Selection line | 3 | -812.95 | 1632.05 | - | - |
| **Leaf Mo** (Al stress; fig. S2) | mm1 | Leaf Mo ~ Treat \| random = Selection line | 7 | -66.94 | 148.50 | -1.73 | 0.0369 $^{***}$ |
| | null | Leaf Mo ~ 1 \| random = Selection line | 3 | -72.05 | 150.23 | - | - |
| **Leaf Mo** (Na stress; fig. S2) | mm1 | Leaf Mo ~ Treat \| random = Selection line | 7 | -133.13 | 280.92 | -43.78 | <0.001 $^{***}$ |
| | null | Leaf Mo ~ 1 \| random = Selection line | 3 | -159.28 | 324.70 | - | - |

**Table S6. Linear mixed-effect models for the analysis of leaf trace-element** (Co, As, Se, Rb, Sr, and Cd; Fig. S6) **concentration of plants under each salt stress in our cross-fostering experiment.** Data analysed, model identification and rationale, number of parameters (*K*), log likelihood (log L), Akaike Information Criterion corrected for finite sample sizes (AICc), difference between selected and null model`s AICc (ΔAICc), and *p*-value. The significantly different values (ANOVA) are indicated: $^{***}$, significant ($p < 0.05$); and $^{ns}$, non-significant. The identification after model ID (e.g., *_vs1*) refers to the variance structure applied to the model in order to satisfy the prerequisites of heteroscedasticity and normality of residues. Using the *nlme* package in R, we used: *_vs1*, varIdent(~1|Treat); *_vs2*, varPower(); *_vs3*, varPower(~fitted(.)|Treat); *_vs4*, varConstPower(); *_vs5*, varComb(varIdent(~1|Treat), varPower()); *_vs6*, varExp(); and *_vs7*, varExp(~fitted(.)|Treat).

| Data | Model ID | Model | *K* | Log L | AICc | ΔAICc | *p* |
|---|---|---|---|---|---|---|---|
| **Leaf Co** (Al stress; fig. S3) | mm1_vs1 | Leaf Co ~ Treat \| random = Selection line | 11 | 927.90 | -1832.29 | -14.03 | <0.001 $^{***}$ |
| | null | Leaf Co ~ 1 \| random = Selection line | 3 | 912.19 | -1818.26 | - | - |
| **Leaf Co** (Na stress; fig. S3) | mm1 | Leaf Co ~ Treat \| random = Selection line | 7 | 751.66 | -1488.64 | -9.98 | 0.001 $^{***}$ |
| | null | Leaf Co ~ 1 \| random = Selection line | 3 | 742.40 | -1478.66 | - | - |
| **Leaf As** (Al stress; fig. S3) | mm1_vs2 | Leaf As ~ Treat \| random = Selection line | 8 | -196.84 | 410.50 | -63.43 | <0.001 $^{***}$ |
| | null | Leaf As ~ 1 \| random = Selection line | 3 | -233.90 | 473.93 | - | - |
| **Leaf As** (Na stress; fig. S3) | mm1_vs6 | Leaf As ~ Treat \| random = Selection line | 8 | -266.81 | 550.48 | -15.44 | <0.001 $^{***}$ |
| | null | Leaf As ~ 1 \| random = Selection line | 3 | -279.89 | 565.92 | - | - |
| **Leaf Se** (Al stress; fig. S3) | mm1 | Leaf Se ~ Treat \| random = Selection line | 7 | 94.70 | -174.78 | -17.84 | <0.001 $^{***}$ |
| | null | Leaf Se ~ 1 \| random = Selection line | 3 | 81.54 | -156.94 | - | - |
| **Leaf Se** (Na stress; fig. S3) | mm1_vs1 | Leaf Se ~ Treat \| random = Selection line | 11 | -74.47 | 172.55 | -55.34 | <0.001 $^{***}$ |
| | null | Leaf Se ~ 1 \| random = Selection line | 3 | -110.88 | 227.89 | - | - |
| **Leaf Rb** (Al stress; fig. S3) | mm1 | Leaf Rb ~ Treat \| random = Selection line | 7 | -398.15 | 810.92 | -13.74 | <0.001 $^{***}$ |
| | null | Leaf Rb ~ 1 \| random = Selection line | 3 | -409.27 | 824.66 | - | - |
| **Leaf Rb** (Na stress; fig. S3) | mm1 | Leaf Rb ~ Treat \| random = Selection line | 7 | -531.95 | 1078.58 | 6.18 | 0.6714 $^{ns}$ |
| | null | Leaf Rb ~ 1 \| random = Selection line | 3 | -533.13 | 1072.40 | - | - |
| **Leaf Cd** (Al stress; fig. S3) | mm1 | Leaf Cd ~ Treat \| random = Selection line | 7 | -556.24 | 1127.12 | -26.53 | <0.001 $^{***}$ |
| | null | Leaf Cd ~ 1 \| random = Selection line | 3 | -573.76 | 1153.65 | - | - |
| **Leaf Cd** (Na stress; fig. S3) | mm1 | Leaf Cd ~ Treat \| random = Selection line | 7 | -567.65 | 1149.98 | -26.42 | <0.001 $^{***}$ |
| | null | Leaf Cd ~ 1 \| random = Selection line | 3 | -585.13 | 1176.40 | - | - |
